# Analysis of 100 high coverage genomes from a pedigreed captive baboon colony

**DOI:** 10.1101/506782

**Authors:** Jacqueline A. Robinson, Shifra Birnbaum, Deborah E. Newman, Jeannie Chan, Jeremy P. Glenn, Laura A. Cox, Jeffrey D. Wall

## Abstract

Baboons (genus *Papio*) are broadly studied in the wild and in captivity. They are widely used as a non-human primate model for biomedical studies, and the Southwest National Primate Research Center (SNPRC) at Texas Biomedical Research Institute has maintained a large captive baboon colony for more than 50 years. Unlike other model organisms though, the genomic resources for baboons are severely lacking. This has hindered the progress of studies using baboons as a model for basic biology or human disease. Here, we describe a dataset of 100 high-coverage whole-genome sequences obtained from the mixed colony of olive (*P. anubis*) and yellow (*P. cynocephalus*) baboons housed at the SNPRC. These data provide a comprehensive catalog of common genetic variation in baboons, as well as a fine-scale genetic map. We show how the data can be used to learn about ancestry and admixture, and to correct errors in the colony records. Finally, we investigated the consequences of inbreeding within the SNPRC colony and found clear evidence for increased rates of juvenile mortality and increased homozygosity of putatively deleterious alleles in inbred individuals.

## INTRODUCTION

Baboons (genus *Papio*) are Old World monkeys commonly found in open woodlands and savannahs across sub-Saharan Africa and the southern portion of the Arabian Peninsula. They are closely related to humans, with an estimated divergence time of ~30 million years (Perelman et al. 2011; Zinner et al. 2013; Pozzi et al. 2014), and like humans, baboons are social, omnivorous, and highly adaptable. They originated in southern Africa and over the past two million years have expanded their range and evolved into six distinct morphotypes, or species: olive (*P. anubis*), yellow (*P. cynocephalus*), hamadryas (*P. hamadryas*), Guinea (*P. papio*), chacma (*P. ursinus*) and kinda (*P. kindae*) baboons (Jolly 1993; Newman et al. 2004; Zinner et al. 2009; Boissinot et al. 2014). Because they share so many similarities with humans, baboons have been studied intensively both in the wild and in captivity since the early 1960s and are considered a useful model in a wide array of research areas.

Over the past several decades, the baboon has become an important non-human primate model in biomedical research, second only to macaques (genus *Macaca*) (reviewed in VandeBerg et al. 2009). Baboons have been used to study normal physiology and development as well as various diseases that commonly affect humans, including diabetes, atherosclerosis, osteoporosis, obesity, hypertension, epilepsy, and addiction (e.g., VandeBerg et al. 2009; Mahaney et al. 2018; Guardado-Mendoza et al. 2009; Aufdemorte et al. 1993; Comuzzie et al. 2003; Szabó et al. 2012). Much of this research has been facilitated by the Southwest National Primate Research Center (SNPRC), which houses the world’s largest captive baboon colony, containing >1,000 individuals at any given time. The colony was established in the 1960s with olive and yellow baboon founders from southern Kenya. Since then, a complete pedigree for the colony spanning seven generations has been maintained, providing relatedness and ancestry information for all captive-bred individuals, along with recorded birth and death dates. Biological samples and phenotype data have also been kept as a resource for biomedical studies.

Relative to other model organisms, however, there are few genomic resources available for baboons. These resources are essential for modern evolutionary and biomedical studies, and their absence hinders baboons’ usefulness as a model organism. The only published baboon genome exclusively used Illumina sequencing (Wall et al. 2016), is highly fragmented, and was assembled into chromosomes based on synteny with the rhesus macaque (*Macaca mulatta*) genome. While there is an olive baboon genome assembly available from NCBI/Ensembl (Panu_3.0), this assembly is also reference-guided and of somewhat poor quality. Further, this ‘public’ baboon assembly has not been freely available to use in scientific publications for more than a decade due to the current interpretation of the Fort Lauderdale Agreement. Very little is known about baboon genomic variation of single nucleotide polymorphisms (SNPs) or haplotypes, and the latest baboon genetic map is based on only 284 microsatellite markers (Cox et al. 2006).

In this study, we take a step towards developing baboon genomic resources by generating high-coverage whole-genome sequence data from 100 SNPRC baboons. We used these genomes to generate a fine-scale map of recombination rates (relative to the Panu_2.0 assembly), for use as a resource in future studies, and to highlight potentially misassembled regions of the reference genome. We also estimated olive vs. yellow baboon ancestry in the sequenced individuals and used this to identify errors in the pedigree file. Lastly, we examined rates of infant mortality and patterns of putatively deleterious variation to investigate the consequences of inbreeding in the colony. We expect the resources and results of this study will provide a useful foundation for future studies of baboons in captivity and in the wild.

## RESULTS

### Resources to enable future baboon genomic research

We sequenced the complete genomes of 100 baboons from the SNPRC colony, including 33 founders, and mapped reads to the Panu_2.0 genome assembly (Supplemental Table S1). In total, we identified 56.4 million SNPs, 5.87 million indels and 5.52 million other complex variants. The raw density of variants was noticeably higher on unplaced scaffolds than on the chromosomes (74.6 variants/kb on scaffolds versus 19.1 variants/kb on autosomes), implying a higher error rate in regions that have not been incorporated into chromosomal assemblies, perhaps due to repetitive sequence content. We focused solely on high quality biallelic SNPs located on the autosomes for our analyses. After applying quality filters (see Methods), our dataset contained 20,352,729 variants distributed across 20 autosomes. Of these, >10.5 million variants were common (minor allele frequency > 0.05) within the sequenced baboon founders, and >653,000 SNPs were highly differentiated (F_ST_ > 0.8) between genetically identified olive and yellow baboon founders (see below). Variation within individuals was comparable to what was observed previously (Wall et al. 2016), with per-individual heterozygosity values ranging from 1.16 to 3.03 heterozygous genotypes per kilobase (kb).

Additionally, we used the variation present within 24 olive founders to generate a fine-scale linkage-disequilibrium based genetic map of the baboon genome with LDhelmet (Chan et al. 2012). Our estimates of recombination rates are in terms of ρ (= 4N_e_r), the population-scaled recombination rate (see Methods). Since LDhelmet is sensitive to a parameter called the “block penalty,” which is a smoothing parameter, we repeated the analysis with a range of values. Our results were largely consistent across block penalty values of 5, 25, and 50 (Supplemental Fig. S1). Here, we focus on results obtained with a block penalty of 5, which the manual suggests is the appropriate value for humans (versus 50 for *Drosophila*). We calculated ρ/bp in nonoverlapping 100 kb windows across the genome and found that estimated rates varied widely, from a low of 3.17 x 10^−7^ to a high of 1.45 ρ/bp within each window. The genome-wide average ρ is ~3.55 per kilobase. We noted that there were several distinct peak regions with exceptionally high recombination rates (Fig. 1A). In total, we identified 45 different 100 kb windows with ρ/bp greater than 100-fold above the mean. Such high recombination rates are biologically implausible and are most likely due to errors in haplotype phasing, reference genome assembly structure, or in the rate estimation itself.

**Figure 1.**
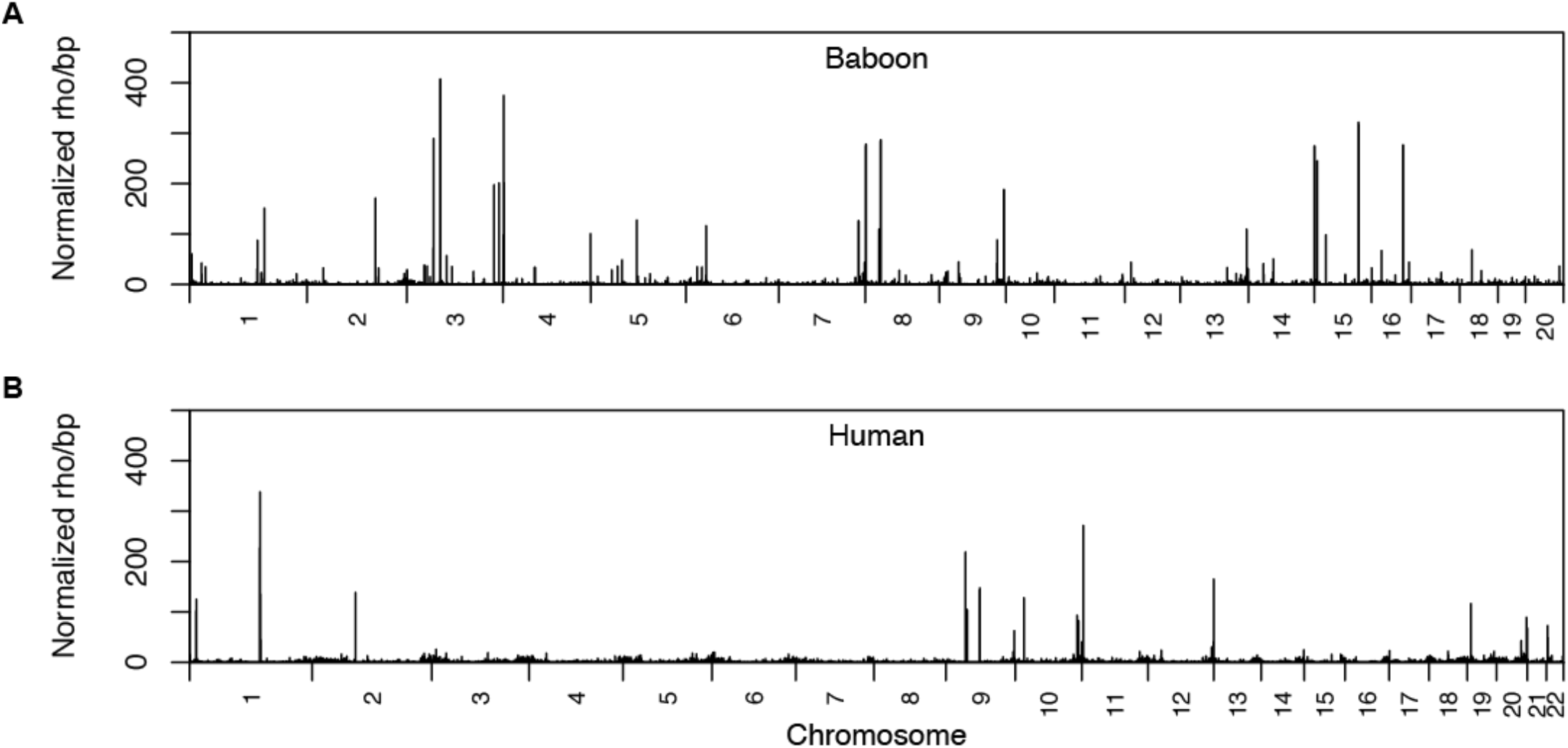
Recombination rates across the baboon and human reference genomes. Rates were inferred from genetic variation in 24 unadmixed olive founder baboons (A) and 24 unrelated African (Yoruban) individuals (B). Rates were calculated in non-overlapping 100 kb windows across the genome and normalized by dividing raw rates by the mean rate inferred within each dataset (mean ρ/bp: baboons, 3.55 x 10’^3^; humans: 5.87 x 10’^3^). Here, a block penalty of 5 was used. Extremely high recombination rates, evident in the large number of high peaks across the genome, highlight putative errors in the Panu_2.0 genome assembly.

To determine whether excessive peaks of recombination rate are expected when the reference genome is well-assembled, we repeated our analysis with a comparable dataset of 24 African (Yoruban) genomes (Wall et al. 2018) (Fig. 1B, Supplemental Fig. S2). We found only 21 different 100 kb windows with ρ/bp greater than 100 times the mean (note that the human genome assembly is slightly longer than the baboon assembly and contains 8.45% more windows overall). The number of windows with exceptionally high ρ in the baboon dataset is significantly greater than in the human dataset (*p*= 1.70 x 10^−14^). Some regions with exceptionally high estimated recombination rates might reflect structural errors in the baboon reference genome assembly (e.g., due to chromosomal rearrangements or structural variants fixed between baboons and rhesus) rather than true biological signal. For example, out of 20 large syntenic differences between a *de novo* Hi-C based baboon assembly and Panu_2.0, 11 show an extremely strong recombination hotspot (corresponding to a complete breakdown of linkage disequilibrium) near at least one of the breakpoints (Batra et al., unpublished data).

### Analysis of founder origins and admixture status

Our data allow us to determine whether species designations in the pedigree were concordant with genetic assignments, and to determine whether any of the founders showed evidence of hybrid origin. The original founders of the SNPRC colony were captured near what is now known to be a large admixture zone between olive and yellow baboons in Eastern Africa (Samuels and Altmann 1986; Tung et al. 2008, Charpentier et al. 2012; Wall et al. 2016). Olive and yellow baboons are phenotypically distinct, and all 33 founders in our sample were originally labeled based on sampling location and physical appearance. Principal component analysis (PCA) with these 33 founders revealed two primary groups corresponding to putative olive and yellow baboons, as well as two distinct outlier individuals (ID: 1X0812 and 1X4384), both originally labeled as olive baboons (Fig. 2A). The PCA results are not consistent with a recent olive-yellow hybrid origin for these individuals, since hybrids would be expected to fall between the olive and yellow baboon clusters. Additionally, one unambiguous olive founder was mislabeled as a yellow baboon (ID: 1X3576). Identity-by-state (IBS) clustering showed the same qualitative patterns (Supplemental Fig. S3), and also suggests that one of the two mystery founders (ID: 1X4384) is genetically more similar to yellow baboons. Without additional information, it is unclear whether the two outlier individuals were members of diverged olive/yellow baboon populations, or were from other baboon species entirely.

**Figure 2.**
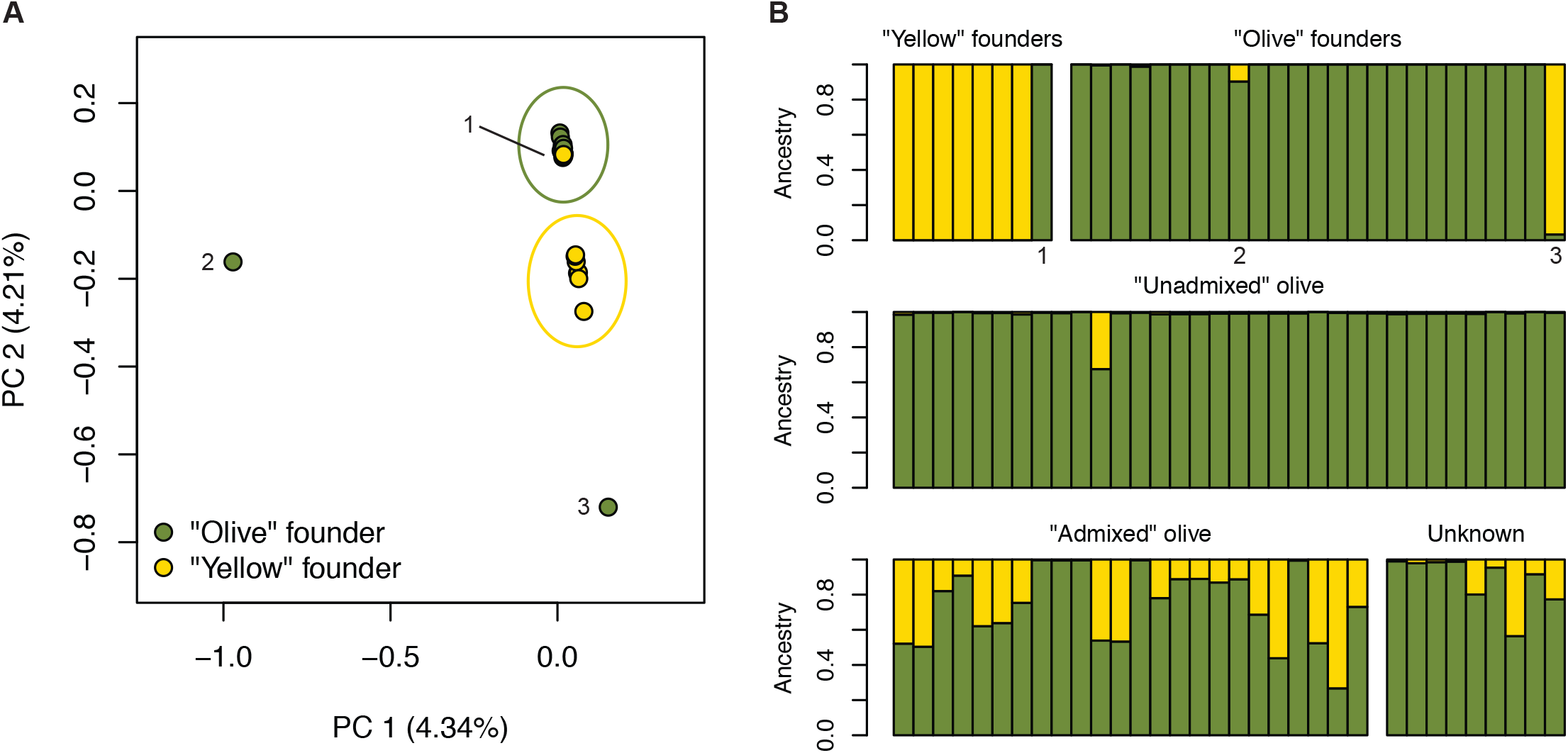
Genetic ancestry patterns in baboons from the SNPRC. *A*) PCA shows that olive and yellow baboon founders form distinct clusters, but one olive baboon was mislabeled as a yellow baboon (1: 1X3576), and two other individuals are extreme outliers (2: 1X0812, 3: 1X4384). *B*) Similarly, ADMIXTURE results under a model of two ancestral populations (K=2) show patterns of olive and yellow baboon ancestry are not always concordant with given species labels. Individuals labeled as 1,2, and 3 in the PCA are indicated. Individuals are grouped according to their classification from the colony records (see Supplemental Table S1).

We ran ADMIXTURE (Alexander et al. 2009) with 31 founders (two mystery individuals were excluded) under a range of K-values (K=1-6) to provide another approach for looking at ancestry and population structure (Supplemental Fig. S4). The most likely number of genetic partitions within the 31 founders was K=2, indicating a clear distinction between 24 genetically olive and 7 genetically yellow baboons and consistent with the PCA and IBS cluster results. Runs with higher K values had higher cross-validation error rates and generated inconsistent partitions within the olive founders. Overall, we found no evidence of recent hybrid ancestry in the founders, and olive and yellow baboons are clearly differentiated. We calculated the genome-wide value of F_ST_ between genetically olive and genetically yellow baboons is 0.366, comparable to previous estimates (Boissinot et al. 2014: F_ST_ =0.3069; Wall et al. 2016: F_ST_ =0.33).

We next used the projection method in ADMIXTURE to assign ancestry proportions to the remaining individuals, assuming two parental populations (K=2) (Fig. 2B). We assigned captive-bred individuals to three groups, according to the pedigree and original species labels of the founders: unadmixed olive individuals, admixed individuals, and individuals with no designation (“unknown”). Of 34 putatively unadmixed captive-born olive baboons, one was a recent hybrid (15197: 32.5% yellow ancestry); of 24 “admixed” individuals, 5 appeared to have pure olive ancestry (>99% olive ancestry); of 9 “unknown” individuals, 4 were clearly admixed (> 5% and <95% olive ancestry). In total, 9 out of 91 individuals (10%) were incorrectly labeled on the basis of species or admixture status. In contrast, the recorded sex of all 100 individuals was correct, which we confirmed by evaluating the ratio of mean coverage on the X chromosome relative to the autosomes (Supplemental Fig. S5). Our results reveal that even in a well-documented captive population with a full pedigree, a non-negligible proportion of individuals may be erroneously categorized. These errors can propagate through the pedigree by affecting the labels of individuals in subsequent generations.

### Comparison of pedigree-based and genomic estimates of inbreeding

A subset of baboons within the SNPRC colony are inbred, making it an ideal system for studying the genomic impact of inbreeding and the link between inbreeding and fitness (i.e. inbreeding depression). The pedigree spans seven generations and contains 16,973 individuals born from 1966-2015. We calculated pedigree-based inbreeding coefficients (F_ped_) for all 16,973 individuals from the pedigree, and found that 1,700 had F_ped_>0 (Fig. 3A). The most common form of inbreeding in the colony is unions between half-siblings or between uncles/aunts and nieces/nephews, resulting in offspring with F_ped_=1/8=0.125 (n=783). The next most common form of inbreeding is mating between parents and offspring (F_ped_=1/4=0.25, n=285). The maximum Fped was 0.40625 (n=3). Thus, the degree of inbreeding within the colony is far greater than what has been observed in human populations (McQuillan et al. 2008; Bittles and Black 2010; Stevens et al. 2012).

**Figure 3.**
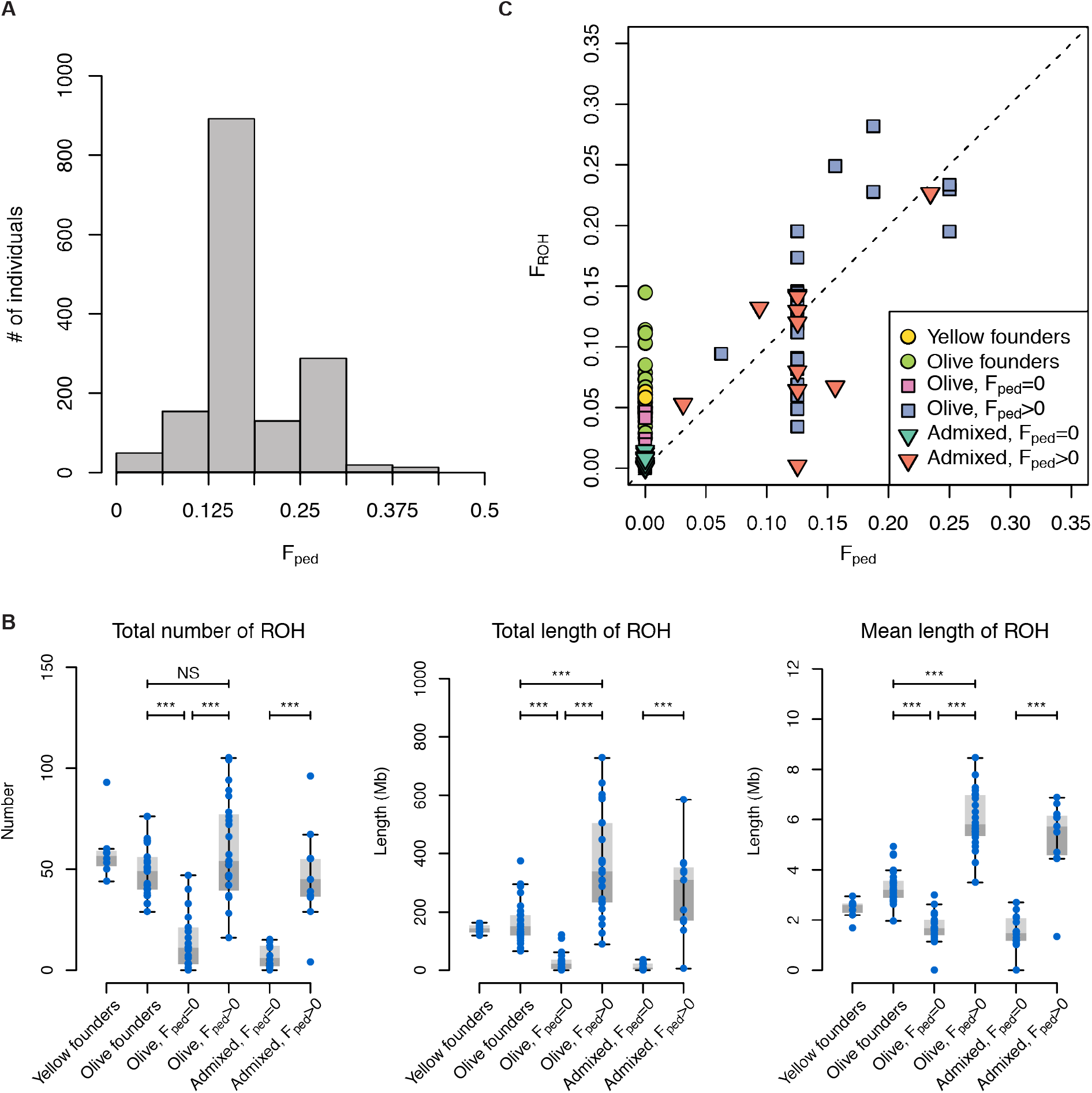
Inbreeding and ROH in the captive baboon colony. *A*) Histogram showing the distribution of inbreeding coefficients in the pedigree (F_ped_). Individuals with F_ped_=0 are not shown. *B-D*) Inbred ancestry (F_ped_ >0) increases the total number, total length, and mean length of ROH in the genome, regardless of admixture status. Founder individuals also show indications of inbreeding in their ancestry. Sample sizes: yellow founders: 8; olive founders: 25; olive, F_ped_=0: 18; olive, F_ped_>0: 24; admixed, F_ped_=0: 14; admixed, F_ped_>0: 11. Significance codes: NS, not significant below a threshold of 0.05; ***, p<0.001. All p-values were multiplied by 4 to correct for multiple tests. *E*) The proportion of the genome contained within ROH (F_ROH_) can vary substantially from F_ped_. The dashed line represents the line *y*=*x*.

Pedigree-based inbreeding coefficients provide the expected values for the proportion of the genome that is autozygous, or inherited identically-by-descent. As expected, we found that inbreeding (Fped>0) is associated with larger numbers of long (>1 Mb) runs of homozygosity (ROH), greater total lengths of ROH in the genome, and longer ROH lengths on average (Fig. 3B). All comparisons between inbred and non-inbred groups among captive-born individuals were statistically significant (*p*≤6.19 x 10^−4^). Interestingly, both olive and yellow baboon founders in our dataset appear to have signatures of inbreeding. Olive baboon founders have greater mean lengths, total numbers, and total lengths of ROH than non-inbred olive baboons born within the colony (*p*≤1.22 x 10^−6^). Long ROH within the founders may reflect non-random mating within wild populations, which have sex-biased dispersal (females are philopatric) and complex social hierarchies that determine access to mates, particularly among males (Alberts et al. 2003).

Next, we compared F_ped_ and F_ROH_, the proportion of the genome within ROH, and found that these values were positively correlated, but F_ROH_ exhibited variance around the predicted values from the pedigree (Fig. 3C). The realized proportion of the genome that is autozygous can vary from the expected value due to inherent randomness in recombination and chromosomal segregation during meiosis. Further, F_ROH_ was greater than F_ped_ in 79% of cases, a statistically significant difference (one-tailed Wilcoxon signed rank test, *p* =6.83 x 10^−7^). This result is consistent with recent studies comparing genomic estimates of inbreeding to pedigree-based estimates, and reflects the fact that genomic data can reveal inbreeding that is not captured in the pedigree (Kardos et al. 2015). Finally, we identified one putatively inbred individual (ID: 8465; F_ped_=0.125) that does not contain autozygous genomic segments consistent with inbred ancestry (F_ROH_=0.00206), suggesting a possible error in the pedigree. Subsequent investigation of animal housing records confirmed that the parentage of this individual listed in the pedigree is incorrect. Overall, our results confirm that genomic measures provide greater resolution and accuracy for quantifying the proportion of the genome that is identical by descent, even relative to estimates from a large pedigree of more than 16,000 individuals over several generations.

### Impacts of inbreeding on juvenile mortality and deleterious variation

To determine whether inbred ancestry is associated with reduced fitness, we calculated the mortality rate of individuals at one day, one week, and one month after birth in 13,313 individuals, 1,393 of which were inbred (Fig. 4A). 16.1% of non-inbred individuals died on their day of birth, versus 23.0% of inbred individuals. This difference corresponds to an odds ratio (OR) for mortality on day of birth of 1.57. The difference in survival between inbred and non-inbred groups was highly statistically significant (*p*=5.24 x 10^−11^). Similar results were obtained for mortality rates within one week and one month of birth. In all comparisons, inbred individuals had significantly higher rates of mortality in early life (*p*≤1.97 x 10^−9^). These results imply that the offspring of related parents have reduced fitness, and suggest that the SNPRC baboon colony would be a useful system for studying the genetic basis of inbreeding depression.

**Figure 4.**
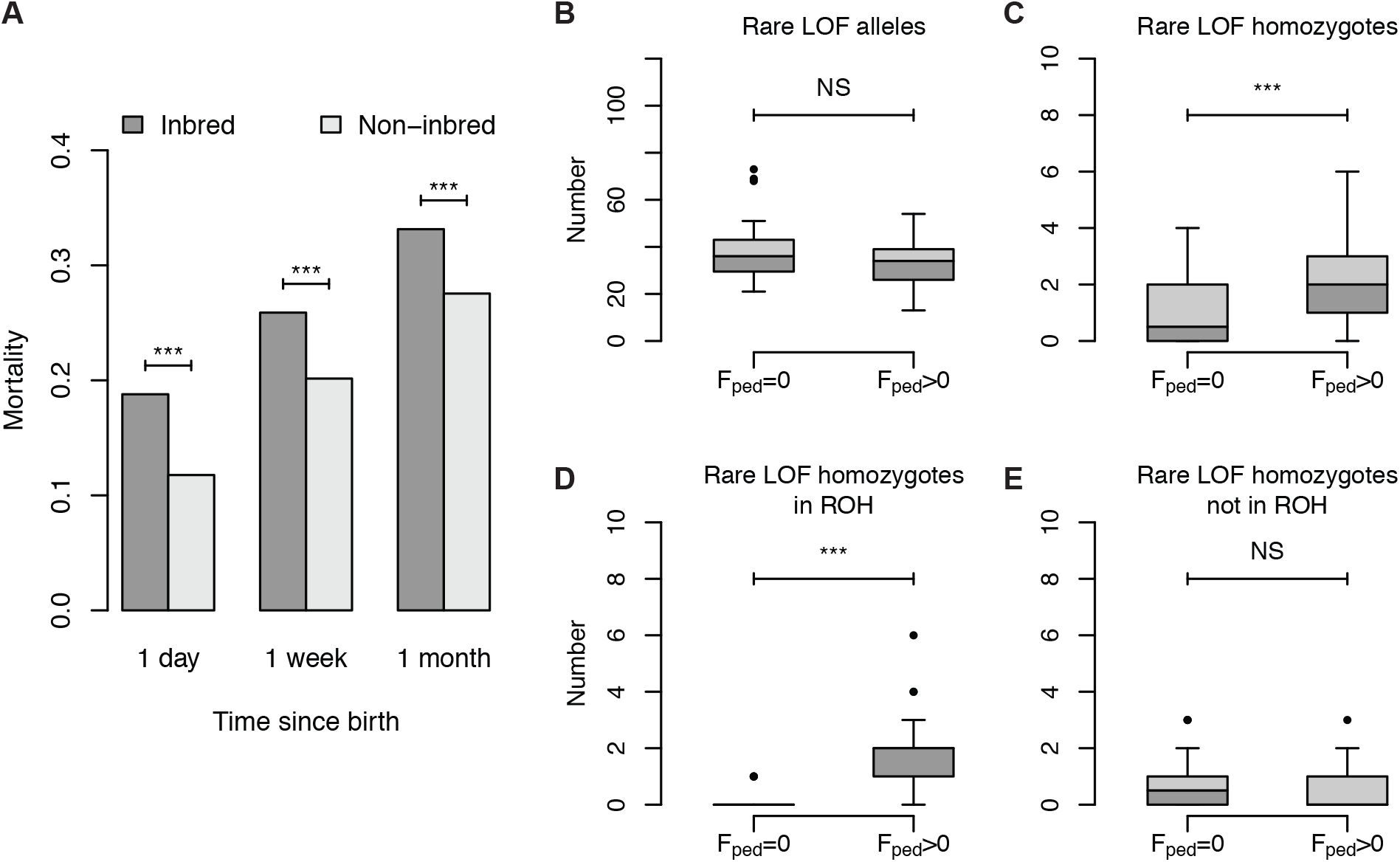
Rates of infant mortality and the burden of LOF mutations as a function of inbreeding in captive-born baboons. *A*) Inbred baboons have substantially higher rates of infant mortality. *B-E*) Boxplots showing the homozygosity of putatively deleterious rare LOF mutations is affected by inbreeding. Specifically, the total number of alleles is not changed by inbreeding, but the number of homozygous LOF mutations increases due to increased ROH content in the genome. The number of rare LOF homozygotes outside of ROH is unchanged between inbred and non-inbred individuals. Significance codes: NS, not significant below a threshold of 0.05; ***, *p*<0.001.

Higher rates of infant mortality in inbred baboons may be due to increased rates of recessive deleterious alleles within ROH. Strongly deleterious recessive alleles can persist in large outbred populations, since these alleles are not exposed to selection when they are rare and most often present in the heterozygous state. Such mutations are often exposed through inbreeding, resulting in inbreeding depression. We compared the total numbers of rare loss-of-function (LOF) alleles and the total numbers of homozygous LOF genotypes between inbred and non-inbred individuals (Fig. 4B, C). LOF mutations diminish or eliminate gene function and are therefore predicted to be deleterious, especially when homozygous and located within genes required for normal function. The total number of LOF alleles was equivalent between inbred and non-inbred groups, consistent with the fact that inbreeding alters genotype frequencies but does not impact allele frequencies in the absence of other factors (e.g. selection, drift). However, inbred individuals had significantly higher numbers of rare homozygous LOF mutations (*p*=1.41 x 10^−4^), specifically due to an increased number of homozygous rare LOF mutations within ROH (*p*= 1.69 x 10^−9^, Fig. 4D). Outside of ROH, rare homozygous LOF mutations were equally prevalent in inbred and non-inbred genomes (Fig. 4E), demonstrating that the increased number of rare homozygous LOF mutations in inbred genomes is due to their higher proportions of ROH, and suggesting a possible role in the reduced fitness of inbred individuals.

## DISCUSSION

The primary goal of our project was to help generate resources that enable future genome-wide studies in baboons, with a focus on the SNPRC baboon colony. To this end, the variants identified in this study could be used for future SNP or capture array designs, while the genetic map will be useful for future evolutionary or genotype-phenotype association studies. In addition, since the method used for estimating recombination rates required phased haplotypes, we also have a computationally phased data set of 24 olive baboon founders that can be used as a reference panel for future genotype imputation in baboons.

Preliminary analyses of our data highlighted several novel findings. First, anomalies in our genetic map identified regions of the Panu_2.0 assembly that might be problematic. Second, genetic analyses identified several errors in the existing animal records related to species identity, admixture status, and inbred ancestry. These inconsistencies may have resulted from mistakes in record-keeping that were never corrected, unexpected matings between baboons in different enclosures, or challenges in distinguishing recently diverged species from one another. Third, we exploited the unique pedigree structure of the colony to quantify the effects of inbreeding on infant mortality and genetic load of rare, homozygous, putatively harmful mutations. Further work on the inbred baboons in the SNPRC colony will provide an ideal opportunity to study the effects of recessive, deleterious mutations in a non-human primate model.

Finally, we want to emphasize that the SNPRC baboon colony, as a mixture of olive and yellow baboons, is also an ideal system for studying the effects of admixture between diverged populations. The yellow and olive baboon founders in our study are highly differentiated (F_ST_ = 0.366), more so than between continental human populations (1000 Genomes Project Consortium 2010) or between isolated human groups (Wall et al. 2008). This differentiation may be useful for the mapping of phenotypic traits (e.g., using admixture mapping), or for evolutionary studies of selection and adaptation. Olive and yellow baboons form a natural hybrid zone in the wild, and the Amboseli Baboon Project has continuously observed several baboon troops in this hybrid zone for over 45 years. We expect that future work enabled by the development of genomic resources will compare the genetic and phenotypic effects of both inbreeding and admixture in the wild and in captivity.

## MATERIALS AND METHODS

### Sequencing and genotype calling

We extracted DNA from archived buffy coats or liver samples from 100 individuals from the SNPRC colony for high coverage (>20X) whole genome sequencing. These individuals included 33 founders of the colony and 67 captive-born descendants (Supplemental Table S1). DNA was extracted using the QIAamp DNA Mini kit (Qiagen) according to manufacturer’s instructions, and DNA quality was assessed using the Qubit BR Assay Kit (Thermo Fisher Scientific) with a Qubit Fluorometer according to manufacturer’s instructions. DNA was quantified using the Kapa Human Genomic DNA Quantification and QC Kit (Kapa Biosystems) according to manufacturer’s instructions. Sequencing libraries were prepared using the TruSeq DNA PCR-Free High Throughput Library Prep Kit (Illumina) according to manufacturer’s instructions with quality assessed using a Bioanalyzer 2100 (Agilent), and quantified using KAPA Library Quantification Kits for Illumina Platform (Kapa Biosystems). Paired-end reads were generated with Illumina HiSeq 2500 technology and then trimmed to remove adapters and low-quality sequence (ea-utils). Trimmed reads were processed with Sentieon Genomics tools (v201611, Freed et al. 2017) to generate variant calls; briefly, reads were aligned to the olive baboon reference genome (Panu_2.0) with BWA MEM (v0.7.12, Li 2013), duplicate reads were removed, and genotypes were called with HaplotypeCaller (McKenna et al. 2010). The mean depth of coverage across individuals was 33.5X. We restricted our analysis to the 21 chromosome-level scaffolds of the reference genome (20 autosomes and one X chromosome), excluding the mitochondrial genome and all unplaced contigs. The X chromosome was used only to calculate the ratio of read depth on the X chromosome versus the autosomes in order to infer each individual’s sex. All other analyses were conducted using only the autosomal data.

We incorporated various filters to minimize the inclusion of erroneous genotypes. We masked repetitive regions using the “Soft-masked” Panu_2.0 reference FASTA available from the UCSC Genome Browser, which annotates repeats based on RepeatMasker and Tandem Repeats Finder (with period of 12 or less) (Smit, 2013; Benson, 1999). Further details are available at https://github.com/priyamoorjani/baboon. We applied recommended hard filters (QD<2.0, FS>60.0, MQ<40.0, MQRankSum<−12.5, ReadPosRankSum<−8.0, S0R>3.0), and excluded variants with excess total depth (DP>4767, the 99^th^ percentile of total depth) or low quality (QUAL<30). Individual genotypes were also filtered to exclude calls with low quality (GQ<20), low coverage (individual read depth <8), or excessive coverage (individual read depth>99^th^ percentile, by individual). Finally, variants that were not biallelic single nucleotide polymorphisms (SNPs), or that had high missingness (>20%) or excess heterozygosity (>50%) were excluded.

### Fine-scale recombination rate estimation

We used LDhelmet (Chan et al. 2012) to infer recombination rates across the genome within 24 unadmixed olive baboon founder genomes and 24 unrelated African (Yoruban) human genomes (GA001430, GA001442, GA001443, GA001444, GA001445, GA001446, GA001447, GA001448, GA001449, GA001450, GA001451, GA001452, GA001453, GA001454, GA001455, GA001456, GA001457, GA001458, GA001459, GA001460, GA001461, GA001462, GA001463, GA000405). The human dataset was derived from a VCF file generated in a separate study (Wall et al. 2018). The human sequence read data are available under NCBI BioProject PRJNA476341.

First, since LDhelmet requires phased input, we phased the genomes using Beagle (v5.0, Browning and Browning 2007). We excluded singletons (of either the reference or alternative allele) since they are uninformative for haplotype phasing, and pruned variants so that no two were within 10 bp of each other, leaving 8,482,860 SNPs in the baboon dataset and 8,505,316 SNPs in the human dataset. We changed the default effective population size parameter to 40,000 for baboons, consistent with the Ne of olive baboons estimated by Boissinot et al. (2014), and 20,000 for humans. Default values were used for all other parameters. Next, we executed the LDhelmet pipeline following the steps outlined in the program manual, using default parameter values except where noted. In particular, we set the population-scaled mutation rate, θ, to 0.0016 in baboons, which is an approximation of the value of θ we calculated from the number of variants in the input data prior to pruning, and 0.001 in humans. We used a window size of 50 and ran the Markov chain Monte Carlo (MCMC) inference for 1 x 10^6^ iterations, following a burn-in period of 1 x 10^5^ iterations. The estimation was conducted under a range of block penalty values (5, 25, 50). Recombination rate estimates are given in units of ρ/bp, where ρ (= 4N_e_*r*) is the population-scaled recombination rate parameter and *r* is the recombination rate per nucleotide per generation. Finally, we converted the recombination rates emitted by LDhelmet, which are given as mean ρ/bp for each SNP interval, into rates in non-overlapping 100 kb windows across the genome. The final window of each chromosome was excluded, since these windows contained fewer sites. A binomial test was used to determine whether there are significantly more windows with extreme recombination rate in the baboon dataset. Here, the number of successes was defined as the number of non-overlapping 100 kb windows in the baboon dataset with estimated recombination rate greater than 100 times the mean (45), the sample size was defined as the number of windows in the baboon genome (25801), and the expected rate of successes was defined as he proportion of high recombination rate windows observed in the human dataset (21/27933).

### Inference of genetic clusters, F_ST_, and admixture

We used SNPRelate (Zheng et al. 2012) to perform PCA and IBS hierarchical clustering, and to calculate a genome-wide value of F_ST_ between olive and yellow baboon founder individuals. Variants within the 33 founder individuals were extracted and pruned for linkage disequilibrium in SNPRelate (threshold=0.2), leaving 127,935 variant sites for PCA and IBS clustering. The IBS clustering analysis consists of constructing a pairwise distance matrix, which is then used to construct a dendrogram to represent genetic similarity between individuals. To estimate genome-wide F_ST_ between olive and yellow baboons, we excluded two founders that were extreme outliers in the PCA (ID: 1X0812 and 1X4384), leaving 17,789,625 unpruned SNPs from the remaining 31 founders. We used VCFtools (Danecek et al. 2011) to calculate F_ST_ on a per-site basis. F_ST_ was calculated using the method of Weir and Cockerham (1984) in both SNPRelate and VCFtools.

We used ADMIXTURE (Alexander et al. 2009) to investigate the possibility of admixed ancestry in the founders and to infer admixture proportions in the captive-born individuals. First, we used the 31 founders that were clearly olive or yellow baboons based on the clustering analyses described above to infer the most likely number of ancestral populations. Here, we ran ADMIXTURE unsupervised under a range of K-values (K=1-6), and then examined the cross-validation errors from each run to determine the most likely K-value. Next, we used the projection method in ADMIXTURE to assign ancestry to the remaining 69 individuals, which is the recommended method for a dataset containing related individuals. The projection analysis consisted of using the population allele frequencies learned from the founders under the most likely K-value to assign ancestry in the remaining individuals.

### Analysis of inbreeding and juvenile mortality in the pedigree

We analyzed a complete pedigree of the baboon colony containing information for individuals born between April 3rd, 1966 and November 4th, 2015. Typically, the sex, birth date, and identity of at least one parent were known for each individual. We used GENLIB (Gauvin et al. 2015) to calculate inbreeding coefficients of all individuals. GENLIB requires that all individuals must be labeled as either male or female, so in cases where the sex of an individual was missing, but the individual was identified as a mother or as a father elsewhere in the pedigree, we filled in the missing sex. In cases where sex was unknown, we arbitrarily changed the sex to female. These individuals have no offspring within the pedigree, therefore their sex is irrelevant for downstream analyses. In general, we defined inbred individuals as any individuals with F_ped_>0, unless otherwise noted.

To compare rates of juvenile mortality in inbred and non-inbred individuals, we calculated the lifespan of all individuals from their recorded birth and “exit” dates. Exit dates can either represent the date an individual died, or the date it left the colony for other reasons. Exit dates within one month of birth were assumed to be due to (natural) mortality. Individuals with no recorded exit date, but that were known to be alive as of November 6^th^, 2015, were given exit dates of November 6^th^, 2015 so that they could be included in analyses of early life survival. Survival rates within one day, one week (7 days), and one month (28 days) of birth were calculated for inbred and non-inbred individuals born between January 1^st^ 1983 and October 7^th^, 2015 (one month before this version of the pedigree was last updated). All individuals born earlier than 1983, which was the first year that any inbred individuals were born, were excluded from mortality analyses. These individuals were excluded since they may have experienced different environmental conditions than inbred individuals, possibly affecting survival in early life. For individuals born late in the pedigree, the one-day, one-week, and one-month survival rates are known with certainty for all individuals born on or before October 7^th^, 2015. Chi-squared tests were used to determine whether rates of juvenile mortality were significantly different between inbred and non-inbred groups.

### Identification of runs of homozygosity

To assess the effects of recent inbreeding in baboon genomes, we used PLINK (v1.9, Chang et al. 2015) to identify large tracts of autozygosity, which indicate genomic regions inherited identically-by-descent from a recent common ancestor shared by both parents. We used the default parameters in PLINK to identify ROH in a pruned set of SNPs as recommended in the manual (*--indep-pairwise 50 5 0.5*). We calculated the proportion of the genome contained within long ROH (1 Mb or longer) as a measure of the realized level of inbreeding (F_ROH_) in each of the sequenced genomes. Here, the numerator for F_ROH_ is the summed length of ROH across the autosomes, divided by the total length of the autosomal genome (2,581,196,250 nucleotides). We used the asymptotic Wilcoxon-Mann-Whitney test from the *coin* library (Hothorn et al. 2008) in R (R Core Team, 2018) to test for significant differences in the total number, total length, and mean length of ROH between groups. Raw *p*-values were multiplied by four to correct for multiple tests (four tests for each ROH statistic evaluated).

### Variant annotation and analysis of LOF mutations

We used SnpEff (Cingolani et al. 2012) to identify the type and impact of mutations, according to the Panu_2.0 genome annotation (Panu_2.0.86). We focused specifically on loss-of-function (LOF) mutations, which are predicted to severely disrupt or eliminate gene function and are therefore most likely to have a negative effect on fitness. This category includes mutations that introduce premature stop codons (“stop_gained”), eliminate start or stop codons (“start_lost”, “stop_lost”), or disrupt splice sites (“splice_acceptor_variant”, “splice_donor_variant”). Only mutations in protein-coding genes were included. Further, we concentrated on LOF mutations that were rare within the founders (<5% frequency), since common LOF mutations are unlikely to be strongly deleterious. We used the asymptotic Wilcoxon-Mann-Whitney test from the *coin* library (Hothorn et al. 2008) in R (R Core Team, 2018) to test for significant differences in the burden of rare LOF mutations between inbred and non-inbred individuals.

## DATA ACCESS

Raw fastq files for all individuals included in this study are freely available from NCBI under BioProject PRJNA433868. A list of accession numbers for each sample is available at https://walllab.ucsf.edu/baboon-data. VCF files and the baboon genetic map generated using LDhelmet, are available at XXX (will be available before publication).

## ACKNOWLEDGMENTS

This work was supported by NIH grants R24 OD017859 (to JDW and LAC) and R01 GM115433 (to JDW). We thank Priya Moorjani for providing the repeat-masked baboon file used in this project.

## SUPPLEMENTAL MATERIAL

**Figure S1.**
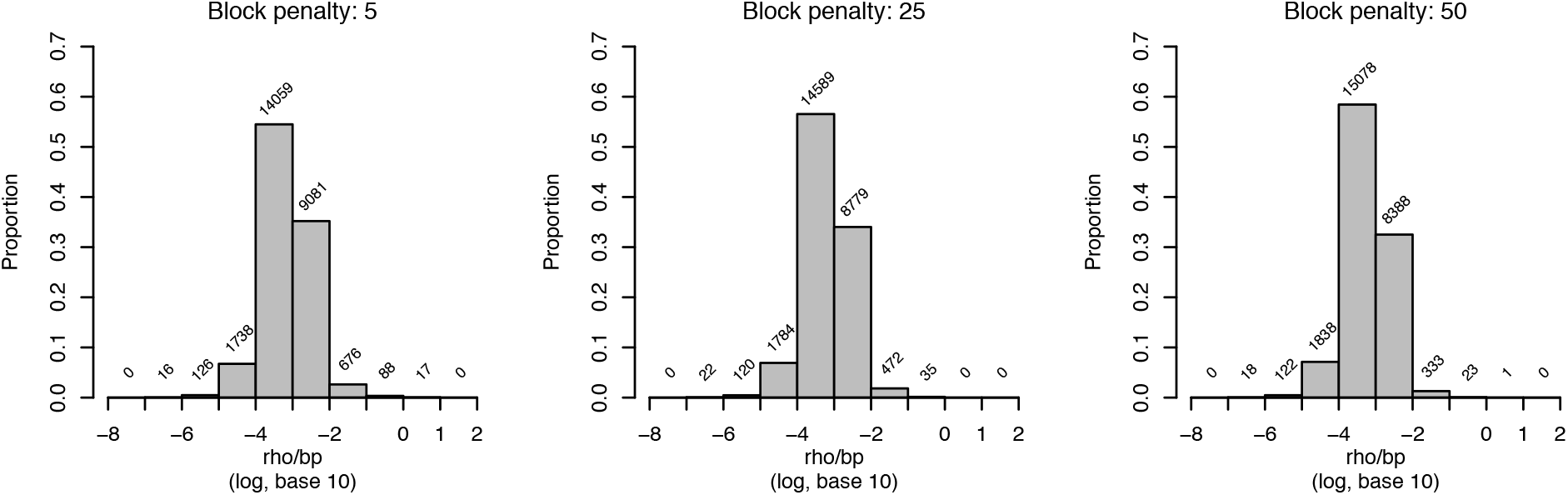
Inferred recombination rates in baboons with varying block penalty values. Estimates of ρ/bp were calculated in non-overlapping 100 kb windows across the genome (see Fig. 1). Numbers above bars indicate the number of windows in each bin. Results are largely consistent across multiple block penalty values.

**Figure S2.**
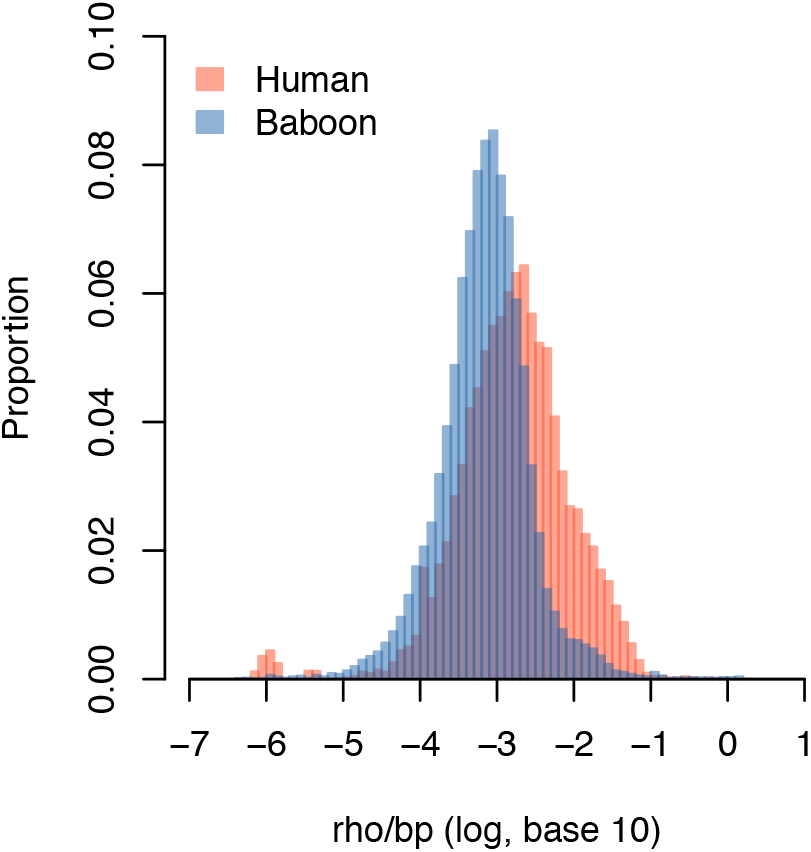
Distribution of inferred recombination rates in humans and baboons. Estimates of ρ/bp were calculated in non-overlapping 100 kb windows across the genome (see Fig. 1). In our analysis, estimates of recombination rates in humans were greater overall than in baboons, but a larger number of windows with extreme ρ values (>100-fold above the mean) were observed in baboons nonetheless.

**Figure S3.**
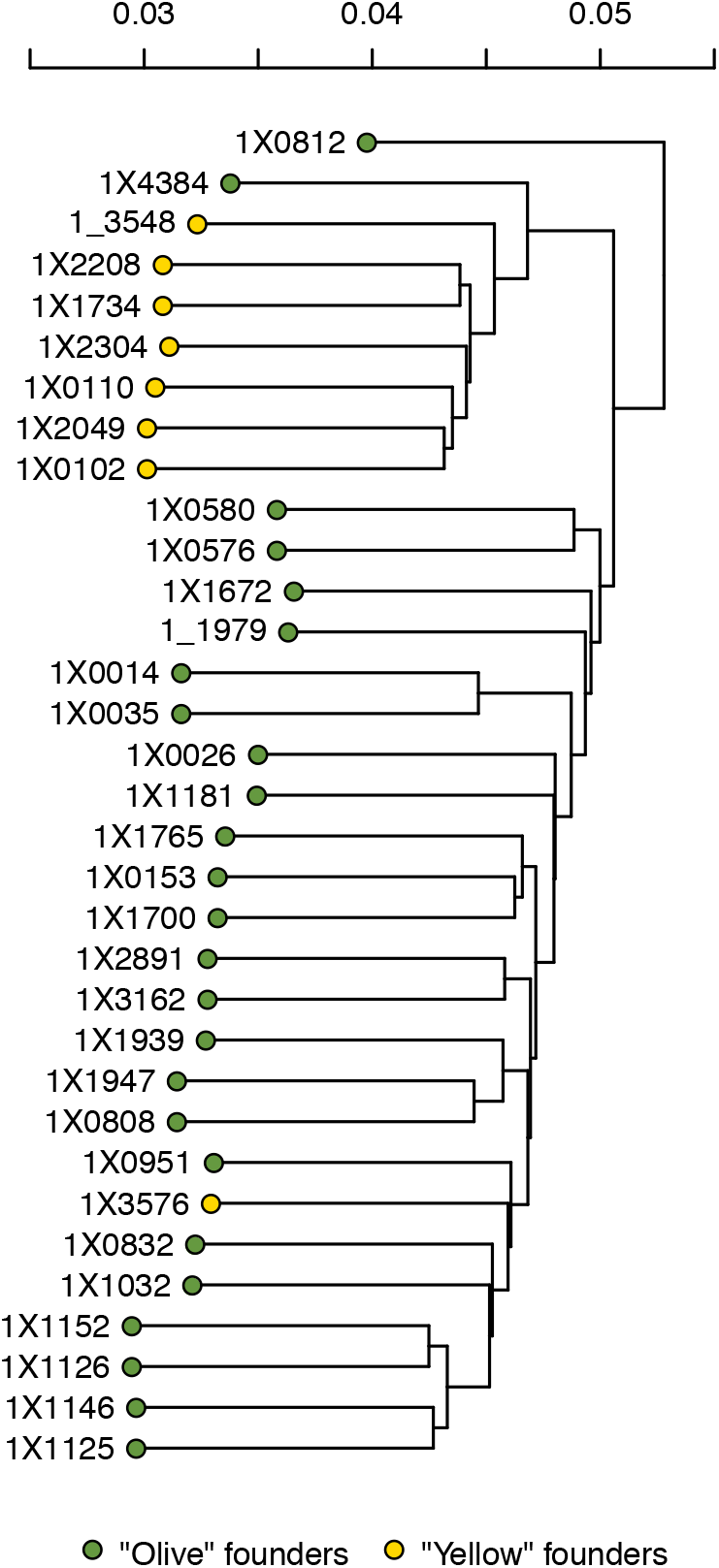
IBS clustering of olive and yellow baboon founders. Identity-by-state analysis shows distinct clusters of olive and yellow baboons, highlighting outliers (1X0812, 1X4384) and a mislabeled sample (1X3576). Results are consistent with patterns from PCA and ADMIXTURE analyses (Fig. 2, Fig. S4).

**Figure S4.**
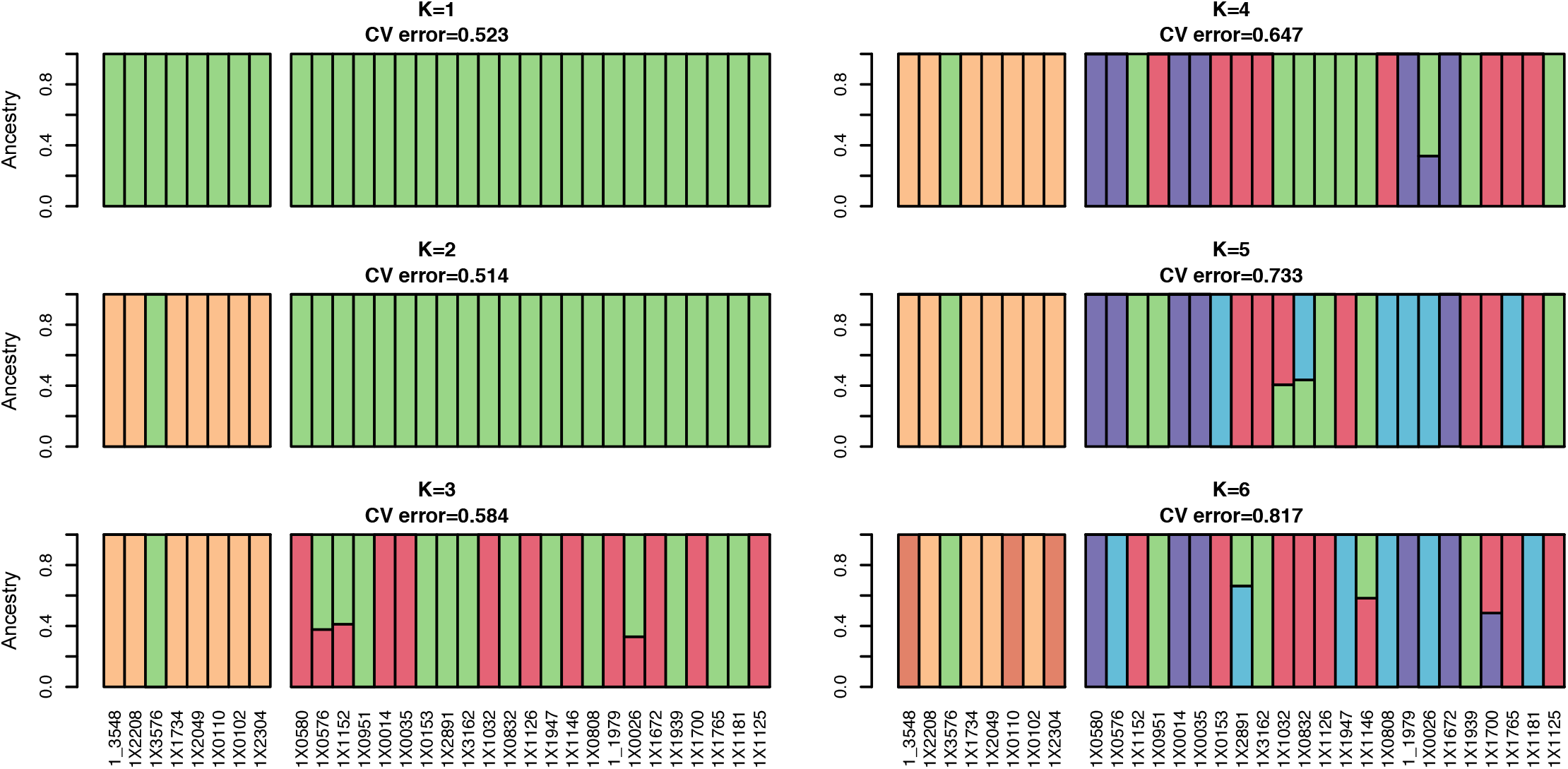
Genetic ancestry of baboon founders under a range of K values. The most likely number of ancestral populations, as indicated by the lowest cross-validation error, in 31 olive and yellow baboon founders is K=2. Here, the extreme outlier founders (1X0812, 1X4384) were excluded. The specimen misidentified as a yellow baboon (1X3576) is apparent, at higher K values, olive and yellow baboon partitions are unchanged, but more partitions within each group appear. None of our results suggest that the founders are the product of recent hybridization between olive and yellow baboons.

**Figure S5.**
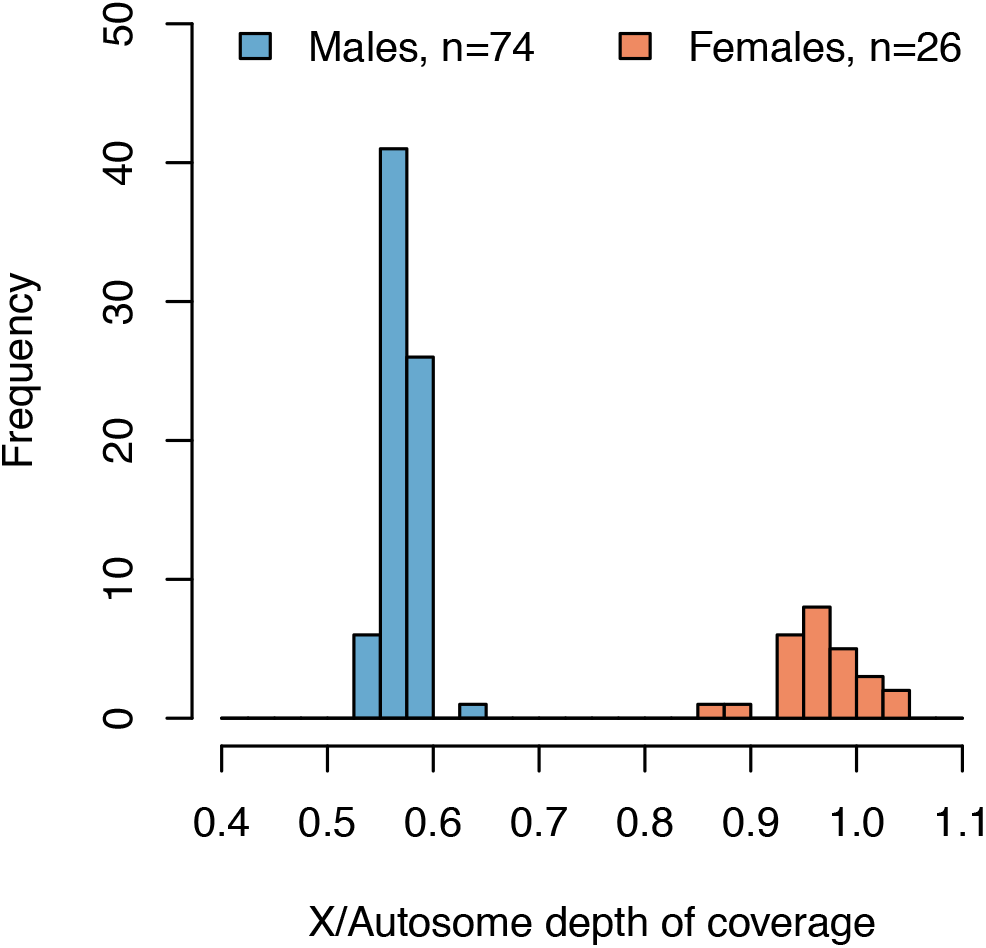
Histogram of the ratio of sequence coverage on the X chromosome relative to the autosomes. Unfiltered depth of coverage values were used. Ratios are consistent with the labeled sex of all individuals, with males having approximately half as much sequence depth on the X chromosome relative to the autosomes.

**Table S1.** List of individuals included in this study. Type, sex, F_ped_ and admixture status are from colony records (e.g. the pedigree).

| Sample ID | Sequencing ID | Type | Sex | $F_{\text{ped}}$ | Admixed |
| --- | --- | --- | --- | --- | --- |
| 1_3548 | FR07886417 | Founder, yellow | F | 0 | No |
| 1X2208 | FR07886421 | Founder, yellow | F | 0 | No |
| 1X3576 | FR07921203 | Founder, yellow | M | 0 | No |
| 1X1734 | FR07921233 | Founder, yellow | F | 0 | No |
| 1X2049 | FR07921237 | Founder, yellow | F | 0 | No |
| 1X0110 | FR07921242 | Founder, yellow | F | 0 | No |
| 1X0102 | FR07921249 | Founder, yellow | M | 0 | No |
| 1X2304 | FR07921251 | Founder, yellow | M | 0 | No |
| 1X0580 | FR07886405 | Founder, olive | F | 0 | No |
| 1X0576 | FR07886409 | Founder, olive | F | 0 | No |
| 1X1152 | FR07886411 | Founder, olive | F | 0 | No |
| 1X0951 | FR07886413 | Founder, olive | M | 0 | No |
| 1X0014 | FR07886416 | Founder, olive | F | 0 | No |
| 1X0812 | FR07886420 | Founder, olive | F | 0 | No |
| 1X0035 | FR07886425 | Founder, olive | F | 0 | No |
| 1X0153 | FR07886426 | Founder, olive | F | 0 | No |
| 1X2891 | FR07919467 | Founder, olive | M | 0 | No |
| 1X4384 | FR07921206 | Founder, olive | M | 0 | No |
| 1X3162 | FR07921212 | Founder, olive | M | 0 | No |
| 1X1032 | FR07921215 | Founder, olive | F | 0 | No |
| 1X0832 | FR07921221 | Founder, olive | M | 0 | No |
| 1X1126 | FR07921226 | Founder, olive | M | 0 | No |
| 1X1947 | FR07921235 | Founder, olive | M | 0 | No |
| 1X1146 | FR07921240 | Founder, olive | F | 0 | No |
| 1X0808 | FR07921241 | Founder, olive | M | 0 | No |
| 1_1979 | FR07921245 | Founder, olive | M | 0 | No |
| 1X0026 | FR07921246 | Founder, olive | F | 0 | No |
| 1X1672 | FR07921258 | Founder, olive | M | 0 | No |
| 1X1939 | FR07921262 | Founder, olive | M | 0 | No |
| 1X1700 | FR07921265 | Founder, olive | M | 0 | No |
| 1X1765 | FR07921266 | Founder, olive | M | 0 | No |
| 1X1181 | FR07921270 | Founder, olive | F | 0 | No |
| 1X1125 | FR07921272 | Founder, olive | F | 0 | No |
| 9841 | 9841 | Captive-born | F | 0 | No |
| 12242 | 12242 | Captive-born | F | 0 | No |
| 15444 | 15444 | Captive-born | F | 12.5 | No |
| 16413 | 16413 | Captive-born | F | 12.5 | No |
| 16517 | 16517 | Captive-born | M | 12.5 | No |
| 17199 | 17199 | Captive-born | F | 12.5 | No |
| 18385 | 18385 | Captive-born | F | 12.5 | No |
| 19181 | 19181 | Captive-born | F | 12.5 | No |
| 19348 | 19348 | Captive-born | F | 12.5 | No |
| 26988 | 26988 | Captive-born | F | 12.5 | No |
| 28246 | 28246 | Captive-born | M | 12.5 | No |
| 1X3837 | FR07886403 | Captive-born | M | 0 | Yes |
| 8653 | FR07886404 | Captive-born | M | 25 | No |
| 1X3697 | FR07886406 | Captive-born | M | 0 | No |
| 15150 | FR07886407 | Captive-born | M | 18.75 | No |
| 15626 | FR07886408 | Captive-born | M | 25 | No |
| 8344 | FR07886410 | Captive-born | M | 0 | Yes |
| 1X3822 | FR07886412 | Captive-born | M | 0 | No |
| 8465 | FR07886414 | Captive-born | M | 12.5 | Unknown |
| 14276 | FR07886415 | Captive-born | M | 3.125 | Yes |
| 7091 | FR07886418 | Captive-born | M | 0 | No |
| 1X2816 | FR07886419 | Captive-born | M | 0 | No |
| 13644 | FR07886422 | Captive-born | M | 0 | Yes |
| 1X4777 | FR07886423 | Captive-born | M | 0 | Yes |
| 7158 | FR07886424 | Captive-born | M | 12.5 | Yes |
| 10192 | FR07918860 | Captive-born | M | 0 | Unknown |
| 8170 | FR07918874 | Captive-born | M | 0 | No |
| 28279 | FR07918876 | Captive-born | M | 12.5 | Yes |
| 28635 | FR07918877 | Captive-born | M | 23.4375 | Yes |
| 15628 | FR07918879 | Captive-born | M | 18.75 | No |
| 9562 | FR07918881 | Captive-born | M | 0 | No |
| 8780 | FR07918882 | Captive-born | M | 0 | No |
| 6716 | FR07918883 | Captive-born | M | 0 | Yes |
| 14930 | FR07918884 | Captive-born | M | 6.25 | No |
| 19207 | FR07919100 | Captive-born | M | 0 | Unknown |
| 14022 | FR07919116 | Captive-born | M | 12.5 | Yes |
| 15156 | FR07919118 | Captive-born | M | 12.5 | No |
| 15197 | FR07919164 | Captive-born | M | 12.5 | Unknown |
| 15633 | FR07919257 | Captive-born | M | 15.625 | Yes |
| 26196 | FR07919258 | Captive-born | M | 0 | Yes |
| 18929 | FR07919259 | Captive-born | M | 0 | Unknown |
| 27351 | FR07919260 | Captive-born | M | 12.5 | Yes |
| 1X4853 | FR07919263 | Captive-born | M | 0 | Yes |
| 9860 | FR07919264 | Captive-born | M | 0 | Yes |
| 14182 | FR07919265 | Captive-born | M | 15.625 | Yes |
| 14925 | FR07919266 | Captive-born | M | 12.5 | No |
| 1X4179 | FR07919267 | Captive-born | M | 0 | No |
| 1X2231 | FR07919268 | Captive-born | M | 0 | No |
| 11887 | FR07919462 | Captive-born | M | 0 | Unknown |
| 15467 | FR07921218 | Captive-born | M | 12.5 | No |
| 14652 | FR07921230 | Captive-born | M | 25 | No |
| 15009 | FR07921250 | Captive-born | M | 12.5 | No |
| 15421 | FR07921261 | Captive-born | M | 12.5 | Yes |
| 15107 | FR07921573 | Captive-born | M | 12.5 | Yes |
| 14712 | FR07921574 | Captive-born | M | 0 | Unknown |
| 14959 | FR07921575 | Captive-born | M | 12.5 | No |
| 15190 | FR07921577 | Captive-born | M | 12.5 | No |
| 13245 | FR07921581 | Captive-born | M | 0 | Unknown |
| 14460 | FR07937948 | Captive-born | M | 9.375 | Yes |
| 10488 | FR07937951 | Captive-born | M | 0 | Yes |
| 10173 | FR07937952 | Captive-born | M | 0 | No |
| 1X4209 | FR07937953 | Captive-born | M | 0 | Unknown |
| 1X3796 | FR07937955 | Captive-born | M | 0 | Yes |
| 10349 | FR07937956 | Captive-born | M | 12.5 | Yes |
| 7937 | FR07937957 | Captive-born | M | 0 | Yes |
| 9656 | FR07937989 | Captive-born | M | 0 | Yes |
| 25347 | FR07937990 | Captive-born | M | 0 | Unknown |

## REFERENCES

1000 Genomes Project Consortium. 2010. A map of human genome variation from population-scale sequencing. Nature 467: 1061.

Alberts SC, Watts HE, Altmann J. 2003. Queuing and queue-jumping: long-term patterns of reproductive skew in male savannah baboons, Papio cynocephalus. Animal Behaviour 65: 821–840.

Alexander DH, Novembre J, Lange K. 2009. Fast model-based estimation of ancestry in unrelated individuals. Genome Research 19: 1655–1664.

Aufdemorte TB, Fox WC, Miller D, Buffum K, Holt GR, Carey KD. 1993. A non-human primate model for the study of osteoporosis and oral bone loss. Bone 14: 581–586.

Batra et al., unpublished data.

Bittles AH and Black ML. 2010. Consanguinity, human evolution, and complex diseases. Proceedings of the National Academy of Sciences 107: 1779–1786.

Boissinot S, Alvarez L, Giraldo-Ramirez J, Tollis M. 2014. Neutral nuclear variation in B aboons (genus P apio) provides insights into their evolutionary and demographic histories. American Journal of Physical Anthropology 155: 621–634.

Browning SR and Browning BL. 2007. Rapid and accurate haplotype phasing and missing data inference for whole genome association studies by use of localized haplotype clustering. American Journal of Human Genetics 81: 1084–1097.

Chan AH, Jenkins PA, Song YS. 2012. Genome-wide fine-scale recombination rate variation in Drosophila melanogaster. PLoS Genetics 8: e1003090.

Chang CC, Chow CC, Tellier LC, Vattikuti S, Purcell SM, Lee JJ. 2015. Second-generation PLINK: rising to the challenge of larger and richer datasets. Gigascience 4: 7.

Charpentier MJ, Fontaine MC, Cherel E, Renoult JP, Jenkins T, Benoit L, Barthes N, Alberts SC, Tung J. 2012. Genetic structure in a dynamic baboon hybrid zone corroborates behavioural observations in a hybrid population. Molecular Ecology 21: 715–731.

Cingolani P, Platts A, Wang le L, Coon M, Nguyen T, Wang L, Land SJ, Lu X, Ruden DM. 2012. A program for annotating and predicting the effects of single nucleotide polymorphisms, SnpEff: SNPs in the genome of Drosophila melanogaster strain w1118; iso-2; iso-3. Fly 6: 80–92.

Comuzzie AG, Cole SA, Martin L, Carey KD, Mahaney MC, Blangero J, VandeBerg JL. 2003. The baboon as a nonhuman primate model for the study of the genetics of obesity. Obesity Research 11: 75–80.

Cox LA, Mahaney MC, VandeBerg JL, Rogers J. 2006. A second-generation genetic linkage map of the baboon (Papio hamadryas) genome. Genomics 88: 274–281.

Danecek P, Auton A, Abecasis G, Albers CA, Banks E, DePristo MA, Handsaker RE, Lunter G, Marth GT, Sherry ST, McVean G. 2011. The variant call format and VCFtools. Bioinformatics 27: 2156–2158.

Freed DN, Aldana R, Weber JA, Edwards JS. 2017. The Sentieon Genomics Tools-A fast and accurate solution to variant calling from next-generation sequence data. bioRxiv doi: 10.1101/115717.

G Benson. 1999. Tandem repeats finder: a program to analyze DNA sequences. Nucleic Acids Research 27: 573–580.

Gauvin H, Lefebvre JF, Moreau C, Lavoie EM, Labuda D, Vézina H, Roy-Gagnon MH. 2015. GENLIB: an R package for the analysis of genealogical data. BMC Bioinformatics 16: 160.

Guardado-Mendoza R, Davalli AM, Chavez AO, Hubbard GB, Dick EJ, Majluf-Cruz A, Tene-Perez CE, Goldschmidt L, Hart J, Perego C, Comuzzie AG. 2009. Pancreatic islet amyloidosis, β-cell apoptosis, and α-cell proliferation are determinants of islet remodeling in type-2 diabetic baboons. Proceedings of the National Academy of Sciences 106: 0906471106.

Hothorn T, Hornik K, van de Wiel MA, Zeileis A. 2008. Implementing a Class of Permutation Tests: The coin Package. Journal of Statistical Software 28: 1–23.

Jolly CJ. Species, subspecies and baboon systematics. 1993. In *Species, Species Concepts, and Primate Evolution* (ed. Kimbel WH, Martin LB). pp. 67–107. Plenum Press, New York.

Kardos M, Luikart G, Allendorf FW. 2015. Measuring individual inbreeding in the age of genomics: marker-based measures are better than pedigrees. Heredity 115: 63.

Li H. 2013. Aligning sequence reads, clone sequences and assembly contigs with BWA MEM. arXiv:1303.3997v2.

Mahaney MC, Karere GM, Rainwater DL, Voruganti VS, Dick EJ Jr, Owston MA, Rice KS, Cox LA, Comuzzie AG, VandeBerg JL. 2018. Diet-induced early-stage atherosclerosis in baboons: Lipoproteins, atherogenesis, and arterial compliance. Journal of Medical Primatology 47: 3–17.

McKenna A, Hanna M, Banks E, Sivachenko A, Cibulskis K, Kernytsky A, Garimella K, Altshuler D, Gabriel S, Daly M, DePristo MA. 2010. The Genome Analysis Toolkit: a MapReduce framework for analyzing next-generation DNA sequencing data. Genome Research 20: 1297–1303.

McQuillan R, Leutenegger AL, Abdel-Rahman R, Franklin CS, Pericic M, Barac-Lauc L, Smolej-Narancic N, Janicijevic B, Polasek O, Tenesa A, Macleod AK, Farrington SM, Rudan P, Hayward C, Vitart V, Rudan I, Wild SH, Dunlop MG, Wright AF, Campbell H, Wilson JF. 2008. Runs of homozygosity in European populations. The American Journal of Human Genetics 83: 359–372.

Newman TK, Jolly CJ, Rogers J. 2004. Mitochondrial phylogeny and systematics of baboons (Papio). American Journal of Physical Anthropology 124: 17–27.

Perelman P, Johnson WE, Roos C, Seuánez HN, Horvath JE, Moreira MA, Kessing B, Pontius J, Roelke M, Rumpler Y, Schneider MP, Silva A, O’Brien SJ, Pecon-Slattery J. 2011. A molecular phylogeny of living primates. PLoS Genetics 7: e1001342.

Pozzi L, Hodgson JA, Burrell AS, Sterner KN, Raaum RL, Disotell TR. 2014. Primate phylogenetic relationships and divergence dates inferred from complete mitochondrial genomes. Molecular Phylogenetics and Evolution 75: 165–183.

R Core Team. 2018. R: A language and environment for statistical computing. R Foundation for Statistical Computing, Vienna, Austria. https://www.R-project.org/.

Rogers J, Mahaney MC, Witte SM, Nair S, Newman D, Wedel S, Rodriguez LA, Rice KS, Slifer SH, Perelygin A, Slifer M, Palladino-Negro P, Newman T, Chambers K, Joslyn G, Parry P, Morin PA. 2000. A genetic linkage map of the baboon (Papio hamadryas) genome based on human microsatellite polymorphisms. Genomics 67: 237–247.

Samuels A and Altmann J. 1986. Immigration of a Papio anubis Male into a Group of Papio cynocephalus Baboons and Evidence for an anubis-cynocephalus Hybrid Zone in Amboseli, Kenya. International Journal of Primatology 7: 131–138.

Smit AFA, Hubley R, Green P. 2015. RepeatMasker Open-4.0. http://www.repeatmasker.org.

Stevens EL, Heckenberg G, Baugher JD, Roberson ED, Downey TJ, Pevsner J. 2012. Consanguinity in Centre d’Etude du Polymorphisme Humain (CEPH) pedigrees. European Journal of Human Genetics 20: 657.

Szabó CÁ, Knape KD, Leland MM, Cwikla DJ, Williams-Blangero S, Williams JT. 2012. Epidemiology and characterization of seizures in a pedigreed baboon colony. Comparative Medicine 62: 535–538.

Tung J, Charpentier MJ, Garfield DA, Altmann J, Alberts SC. 2008. Genetic evidence reveals temporal change in hybridization patterns in a wild baboon population. Molecular Ecology 17: 1998–2011.

VandeBerg, J. L., S. Williams-Blangero, and S. D. Tardif (eds.). 2009 The baboon in biomedical research. Springer, New York, NY.

Wall JD, Cox MP, Mendez FL, Woerner A, Severson T, Hammer MF. 2008. A novel DNA sequence database for analyzing human demographic history. Genome Research 18: 1354–1361.

Wall JD, Schlebusch SA, Alberts SC, Cox LA, Snyder-Mackler N, Nevonen KA, Carbone L, Tung J. 2016. Genomewide ancestry and divergence patterns from low-coverage sequencing data reveal a complex history of admixture in wild baboons. Molecular Ecology 25: 3469–3483.

Wall JD, Stawiski E, Ratan A, Kim HL, Kim C, Gupta R, Suryamohan K, Gusareva ES, Purbojati RW, Bhangale T, et al. 2018. The GenomeAsia 100K Project: Enabling genetic discoveries across Asia, in revision.

Weir BS and Cockerham CC. 1984. Estimating F-Statistics for the analysis of population structure. Evolution 38: 1358–1370.

Zheng X, Levine D, Shen J, Gogarten SM, Laurie C, Weir BS. 2012. A high-performance computing toolset for relatedness and principal component analysis of SNP data. Bioinformatics 28: 3326–3328.

Zinner D, Groeneveld LF, Keller C, Roos C. 2009. Mitochondrial phylogeography of baboons (Papio spp.) – Indication for introgressive hybridization? BMC Evolutionary Biology 9: 83.

Zinner D, Wertheimer J, Liedigk R, Groeneveld LF, Roos C. 2013. Baboon phylogeny as inferred from complete mitochondrial genomes. American Journal of Physical Anthropology 150:133–140.

